# Coriander leaf essential oil as an immunomodulator: enhancing *NF-κB*-driven RAW246.7 murine macrophages response to *Candida albicans*

**DOI:** 10.1101/2025.11.09.687462

**Authors:** Pratsanee Hiengrach, Pasinee Sangsiwarit, Waewta Kuwatjanakul, Kittipan Samerpitak, Pitchaya Luksanawilas

## Abstract

Coriander is a cultivated aromatic herb that has been used in both culinary and medical treatment worldwide. Coriander essential oil contains various bioactive components that can have antibacterial, antioxidant, and anti-inflammatory effects. In addition, the oil has antifungal activity against *Candida albicans* and suppresses its biofilm formation. Besides, the oil can increase macrophage phagocytic activity. However, its effect on other macrophage functions during *C. albicans* infection is still unclear. This study aims to investigate the effect of coriander leaf essential oil on macrophage activity and inflammatory signaling during *C. albicans* infection. RAW264.7, a murine macrophage cell line, was cultured with viable *C. albicans*, either in the absence or presence of the oil (0-50 μg/mL). The killing activity, pro- /anti-inflammatory cytokine production, and *NF-κB* gene expression were assessed. This study revealed the potential of coriander leaf essential oil as an immunomodulator that enhances macrophage responses to *C. albicans* via the *NF-κB* pathway. These findings may help to further the development of coriander leaf essential oil as an adjuvant antifungal and immunomodulatory medication.

## Introduction

Coriander (*Coriandrum sativum*), also known as Chinese parsley, is an aromatic and therapeutic annual plant belonging to the family Apiaceae (1). It is widely cultivated across diverse climatic regions worldwide. In Thai cuisine, fresh coriander leaf is frequently used as a garnish for rice dishes, soups, and curries. Coriander has been reported to have potential in traditional medicinal therapies of several health conditions, such as anxiety, constipation, diabetes mellitus, dyspepsia, parasitic infection, irritable bowel syndrome (IBS), and dermatological irritation (2, 3). Coriander essential oil, typically extracted from dried seeds or leaves via steam distillation, has been reported to have antimicrobial properties that prevent bacterial contamination in food. Additionally, it contains bioactive compounds with antioxidant and anti-inflammatory effects (4). Previous studies demonstrated that coriander essential oil exerted potent antifungal activity against *Candida* species, particularly *C. albicans* (5), the most common opportunistic fungal pathogen (6, 7). *Candida albicans* can infect multiple sites, such as the oral cavity, skin, genital organs, and bloodstream. The coriander essential oil binds to ergosterol in the fungal cell membrane, leading to increased ionic permeability, membrane disruption, and cell death. Moreover, it could inhibit *Candida* biofilm formation, thereby reducing fungal adherence and pathogenicity (8). Besides, coriander essential oil has been reported ability to modulate host immune responses. It enhanced the phagocytic capacity of macrophages, the key innate immune cells responsible for the early recognition and elimination of fungal pathogens. However, the influence of the oil on other macrophage functions, such as killing activity and cytokine-mediated inflammatory mechanisms during *C. albicans* infection remains unclear. Therefore, this study aims to investigate the effects of coriander essential oil on macrophage function and inflammatory signaling pathways during *C. albicans* infection. The result may offer novel insights into the potential use of coriander essential oil as a natural adjuvant for fungal infection therapy.

## Methods

### Ethical approvement

The research protocol employed in this study was reviewed and approved by the Center for Ethics in Human Research, Khon Kaen University (ECKKU), Thailand. Ethical approval was granted under the reference number HE671565.

### Cultivation of murine macrophage RAW264.7 cells

The RAW264.7, murine macrophage cell line, was maintained in RPMI 1640 medium (Thermo Fisher Scientific, Waltham, MA, USA), supplemented with 10% (v/v) heat-inactivated fetal bovine serum (FBS) (Sigma-Aldrich, St. Louis, MO, USA) and 2 mM L-glutamine. The cells were incubated at 37°C in a humidified atmosphere containing 5% CO_2_.

### Cultivation of *C. albicans*

*C. albicans* strain ATCC90028 (Fisher Scientific, Waltham, MA, USA), obtained from the American Type Culture Collection, was cultured on Sabouraud Dextrose Agar (SDA; Oxoid, Basingstoke, Hampshire, UK) at 37°C for 48 h. Then, the yeast cells were suspended in 1x phosphate-buffered saline (PBS), pH 7.4, and adjusted to a final concentration of 1 × 10^5^ CFU/mL before experimentation.

### Liquid chromatography-mass spectrometry of coriander leaf essential oil

Laboratory-grade coriander leaf essential oil was obtained from Nature in Bottle Co., Ltd. (New Delhi, India). The separation and quantification of its constituents were performed using a liquid chromatography-mass spectrometry (LC–MS) system (U2Bio, Thailand). In brief, 150 µL of coriander leaf essential oil was mixed with 150 µL of 70% methanol (MeOH) containing 100 ng/mL sulfadimethoxine as an internal standard. The mixture was centrifuged at 14,000 rpm for 10 min, and the supernatant was transferred to an LC–MS vial for analysis. Chromatographic separation was carried out using a Poroshell 120 EC-C18 column (2.1 × 100 mm, 2.7 µm) maintained at 50°C. The injection volume was 10 µL. The mobile phases consisted of 0.1% formic acid in water (solvent A) and 0.1% formic acid in acetonitrile (solvent B), with a flow rate of 0.4 mL/min.

### Evaluation of macrophage-associated parameters

#### Measurement of killing activity

To evaluate the candidacidal activity of macrophages, RAW264.7 cells were seeded into a 96-well microplate at a density of 1 × 10^4^ cells/well. Following overnight incubation, the viable

*C. albicans* were co-incubated with the macrophages at 37°C in a 5% CO_2_ atmosphere for 0, 6, and 18 h, either in the absence or presence of coriander essential oil (0-50 μg/mL). After each incubation period, 100 µL of the co-culture was plated onto SDA and incubated at 37°C for 72 h. The *Candida* colonies were subsequently enumerated to assess macrophage-mediated killing activity.

### Measurement of pro- and anti-inflammatory cytokines

To investigate cytokines that are secreted by macrophages. RAW 264.7 cells will be seeded into a 96-well microplate overnight at a density of 1 × 10^4^ cells*/*well. Then, live *Candida* yeast cells will be co-incubated at 37ºC in a 5% CO_2_, incubated for 0, 6, and 18 h, with or without coriander extract oil (0-50 μg/mL). After the challenge, the supernatant will be collected for the determination of the amount of pro-inflammatory cytokines (IL-6 and TNF-α) and anti-inflammatory cytokines (IL-10) using enzyme-linked immunosorbent assay (ELISA; Invitrogen, Massachusetts, USA). All cytokines will be quantified by a spectrophotometer at an excitation wavelength of 450 nm and an emission at 570 nm.

### Measurement of gene-associated inflammatory pathway

To investigate the inflammatory signaling pathway in RAW264.7 macrophages, the expression of *NF-κB* was evaluated at the transcriptional level using quantitative reverse transcription polymerase chain reaction (qRT-PCR). RAW264.7 cells were seeded into 96-well microplates at a density of 1 × 10^4^ cells per well and incubated overnight. Subsequently, viable *C. albicans* cells were co-incubated with the macrophages at 37°C in a humidified 5% CO_2_, incubated for 0, 6, and 18 h, either in the absence or presence of coriander leaf essential oil (0-50 μg/mL). Macrophages were collected from each condition for RNA extraction using TRIzol reagent (Invitrogen, Massachusetts, USA), following the manufacturer’s protocol (9). Complementary DNA (cDNA) was synthesized using the high-capacity cDNA reverse transcription kit (Thermo Fisher Scientific, Massachusetts, USA). The qRT-PCR was performed using Maxima SYBR Green qPCR Master Mix (Thermo Fisher Scientific) on a CFX96 touch real-time PCR detection system (Bio-Rad, California, US). Gene expression levels were analyzed using the ΔCt method, following the previous publication (10). The following primers were used for *NF-κB*: forward, 5’-AGCCAGCTTCCGTGTTTGTT-3’ and reverse, 5’-AGGGTTTCGGTTCACTAGTTTCC-3’ (10). Experiments were repeated three times.

## Results

### Compound components of coriander leaf essential oil

The coriander leaf essential oil has been identified as a rich source of diverse phytochemical compounds, both identified and unidentified compounds. The identified compound, including various mono-, di-, and triterpenoids, fatty acids, and naphthofuran derivatives (Table 1). Among the monoterpenoids that were detected are thymol, carveol, and chrysanthemic acid. Besides, an essential oil also contained several di- and triterpenoids, such as darutigenol, lagochilin, Amyrin, darutigenol, sterebin E, sterebin G, and abietic acid, along with a range of labdane- and abietane-type diterpenoids. In addition to terpenoids, coriander leaf essential oil was found to contain a complex mixture of fatty acids. Saturated fatty acids, such as lauric acid, capric acid, undecanoic acid, dodecanoic acid, decanoic acid, and hydroxydodecanoic acid, were identified. Moreover, several bioactive oxylipins, including 3-Oxo-Dodecanoic acid, 11-hydroxyeicosatetraenoic acid (11-HETE), 9-hydroxyeicosapentaenoic acid (9-HEPE), and 12-oxo-eicosatetraenoic acid (12-OxoETE), were detected (Figure 1A-B).

**Table 1.**
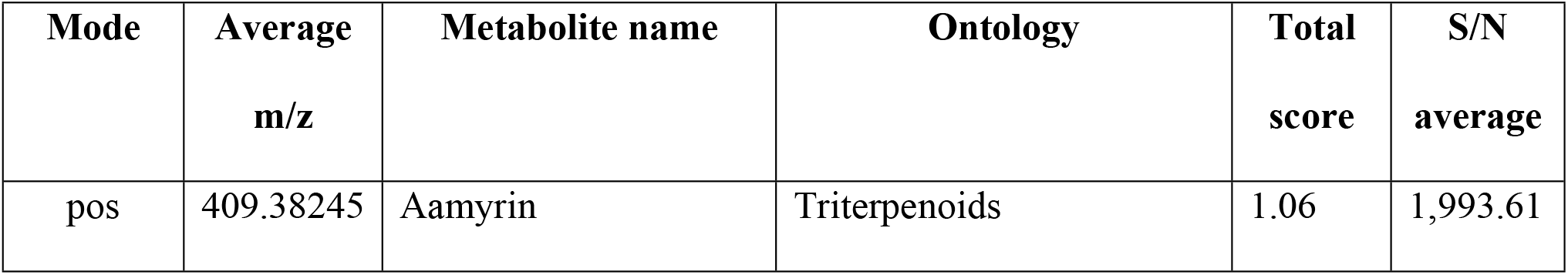
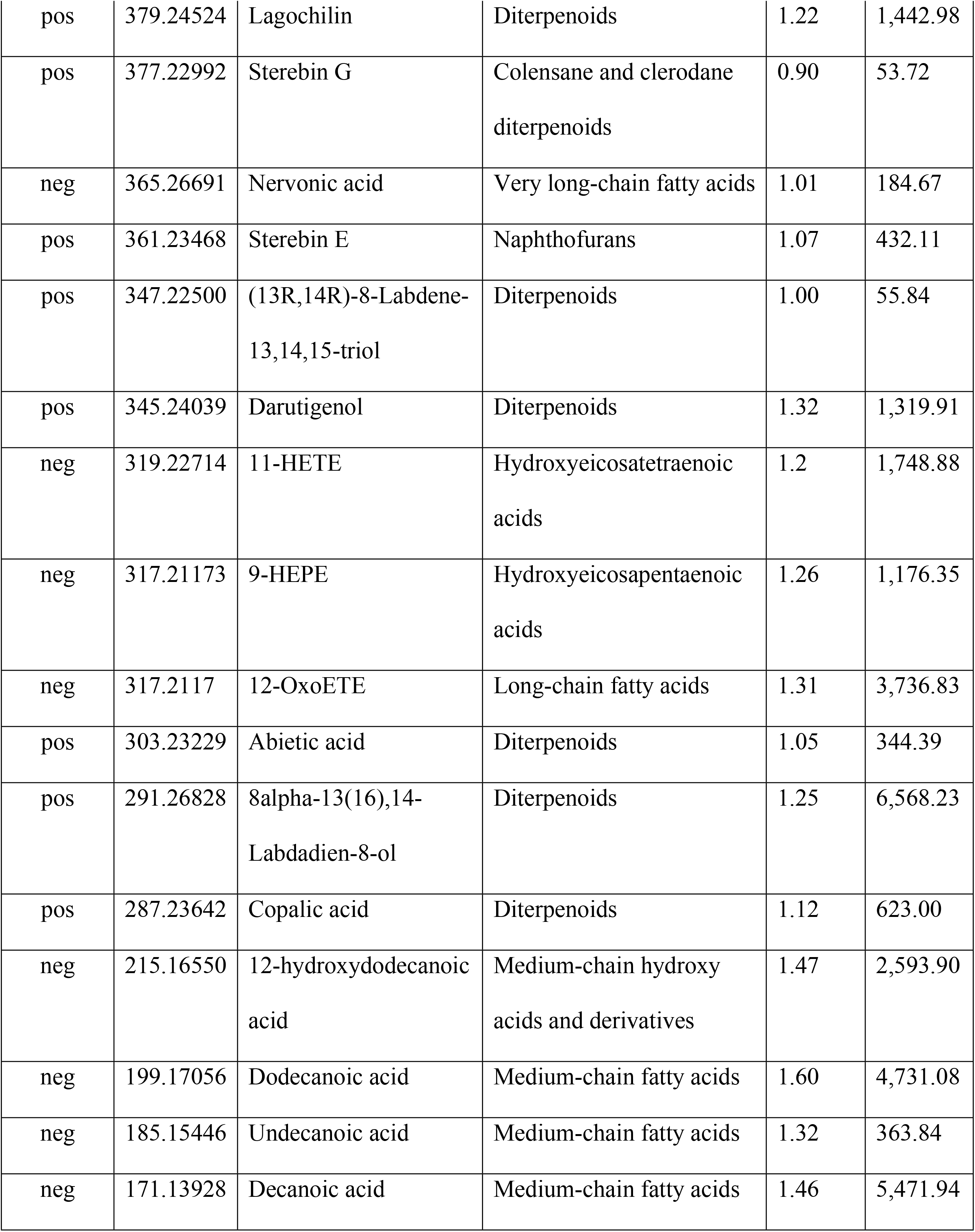
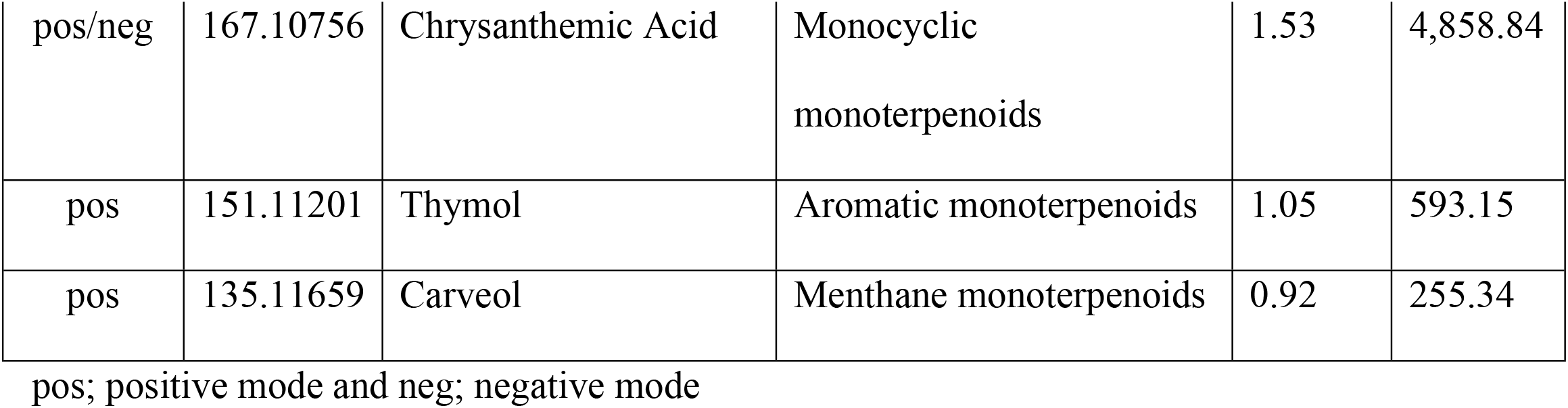
Phytochemical profile of coriander leaf essential oil using the LC-MS method.

**Figure 1.**
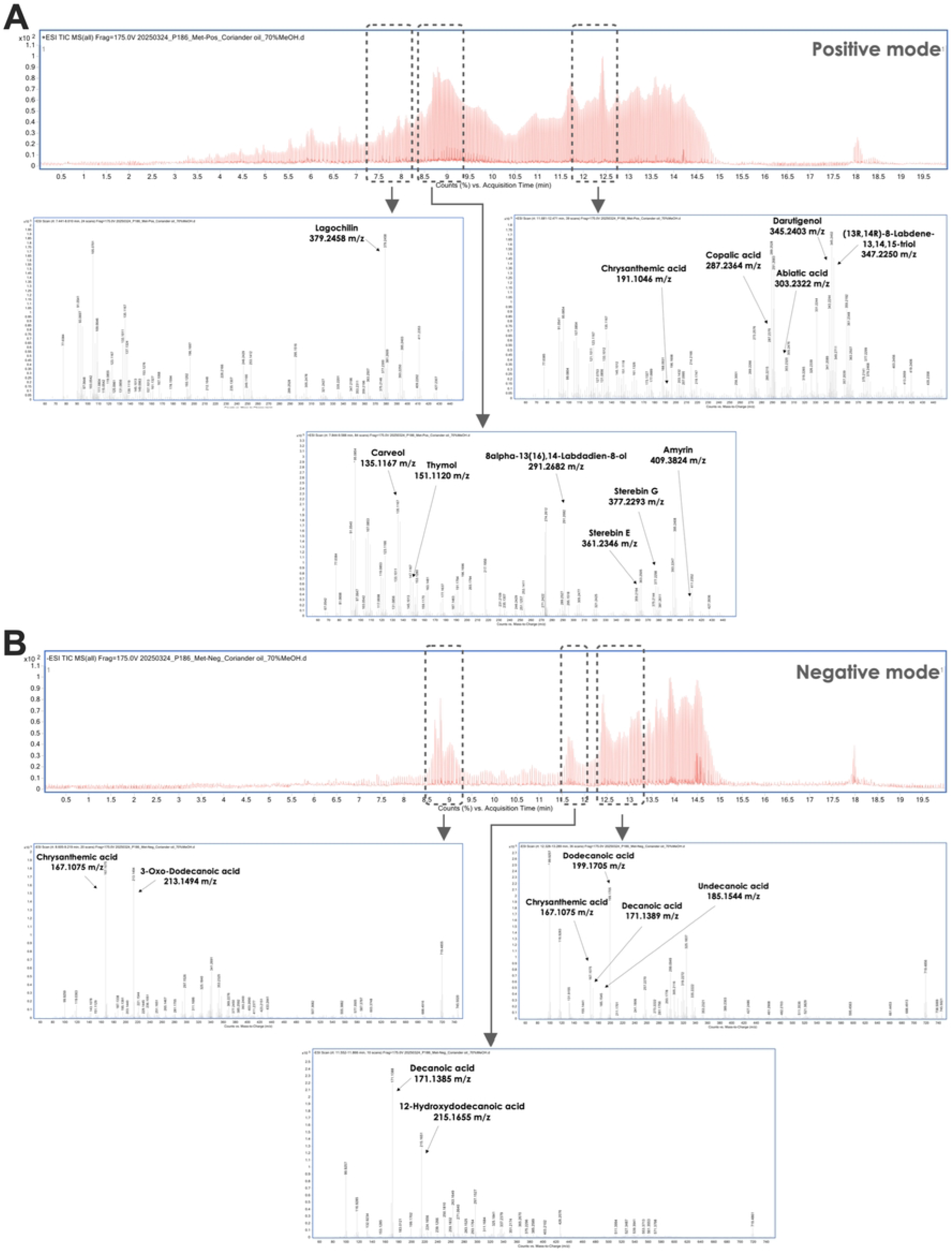
A chromatography of coriander leaf essential oil analyzed by LC-MS, showing results in positive ion mode (upper panel) and negative ion mode (lower panel).

### The effects of coriander leaf essential oil on macrophage function and inflammatory signaling pathways during *C. albicans* infection

Due to their involvement in host defense mechanisms against fungal pathogens, the ability of macrophages to exert killing activity and cytokine production was evaluated. The results demonstrated that coriander leaf essential oil at concentrations between 5–50 µg/mL significantly reduced the number of *C. albicans* colonies at both 6 and 18 h in comparison with the untreated *C. albicans* control group. Furthermore, when coriander leaf essential oil was administered in conjunction with RAW264.7 cells, a statistically significant reduction in *C. albicans* colony count was observed compared to the untreated co-culture group (RAW264.7 + *C. albicans*). This reduction displayed a dose-dependent trend. Notably, coriander leaf essential oil at a concentration of 50 µg/mL significantly enhanced the killing activity of RAW264.7 cells against *C. albicans* (Figure 2A).

**Figure 2.**
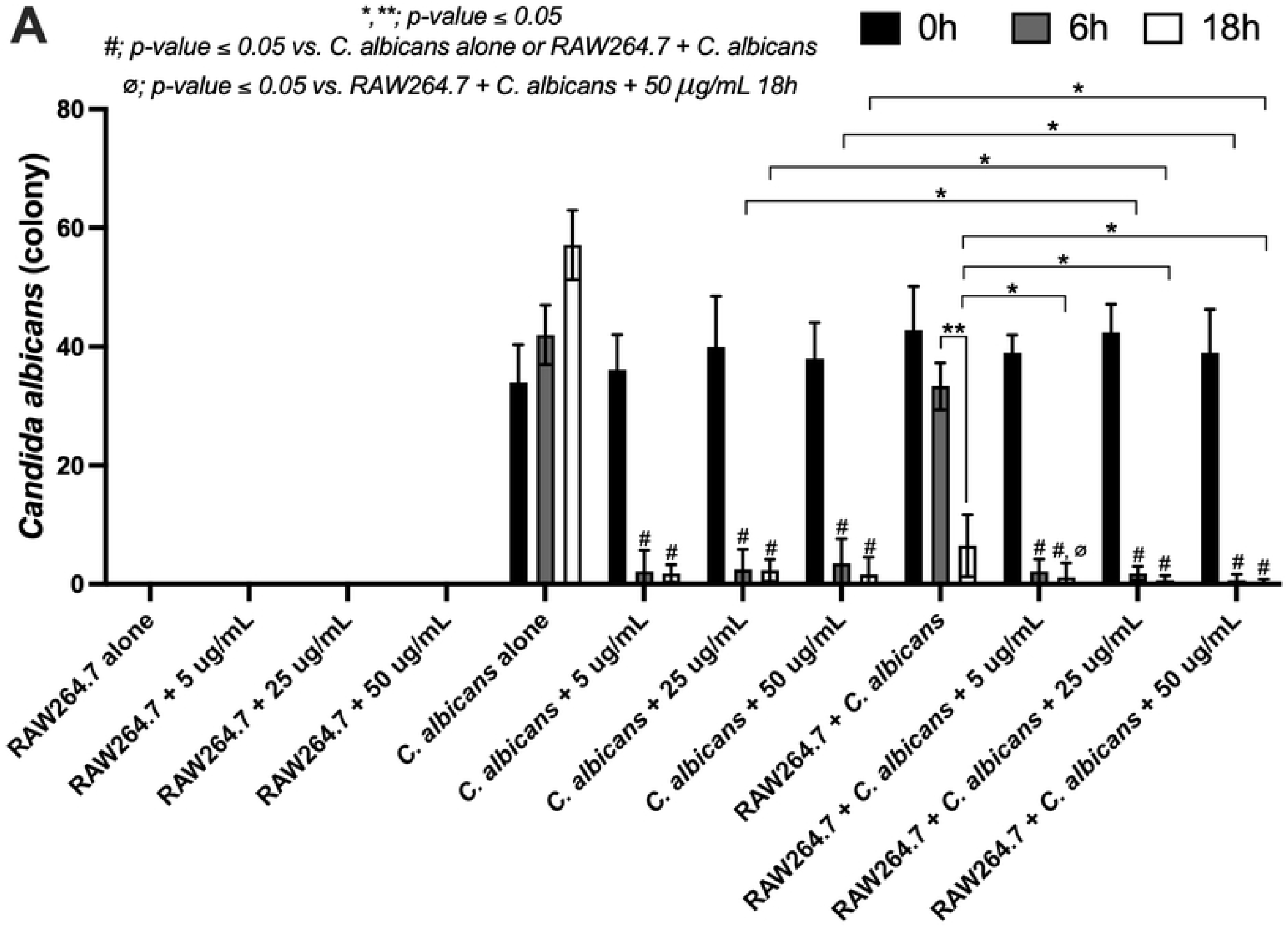
The characteristics of the RAW264.7 macrophage cell line after activation by *C. albicans*, either in the absence or presence of coriander essential oil (0-50 μg/mL), are demonstrated based on killing activity analysis. Data were collected in triplicate. The data are shown as the mean ± SE, *, **; *p-value* ≤ *0*.*05*, #; *p-value* ≤ *0*.*05* vs. *C. albicans* alone or RAW264.7 + *C. albicans*, and ∅; *p-value* ≤ *0*.*05* vs. RAW264.7 + *C. albicans* + 50 μg/mL 18 h, between the indicated groups using ANOVA with Tukey’s analysis.

The production of pro-inflammatory cytokines indicates that coriander leaf essential oil induces an immune response with an additive effect, as evidenced by elevated levels of both TNF-α (Figure 3A) and IL-6 (Figure 3B) compared to other conditions. However, no significant differences were observed across time points or among coriander leaf essential oil concentrations ranging from 5–50 μg/mL. In contrast, the anti-inflammatory cytokine, IL-10, showed no significant differences among the tested conditions (Figure 3C).

**Figure 3.**
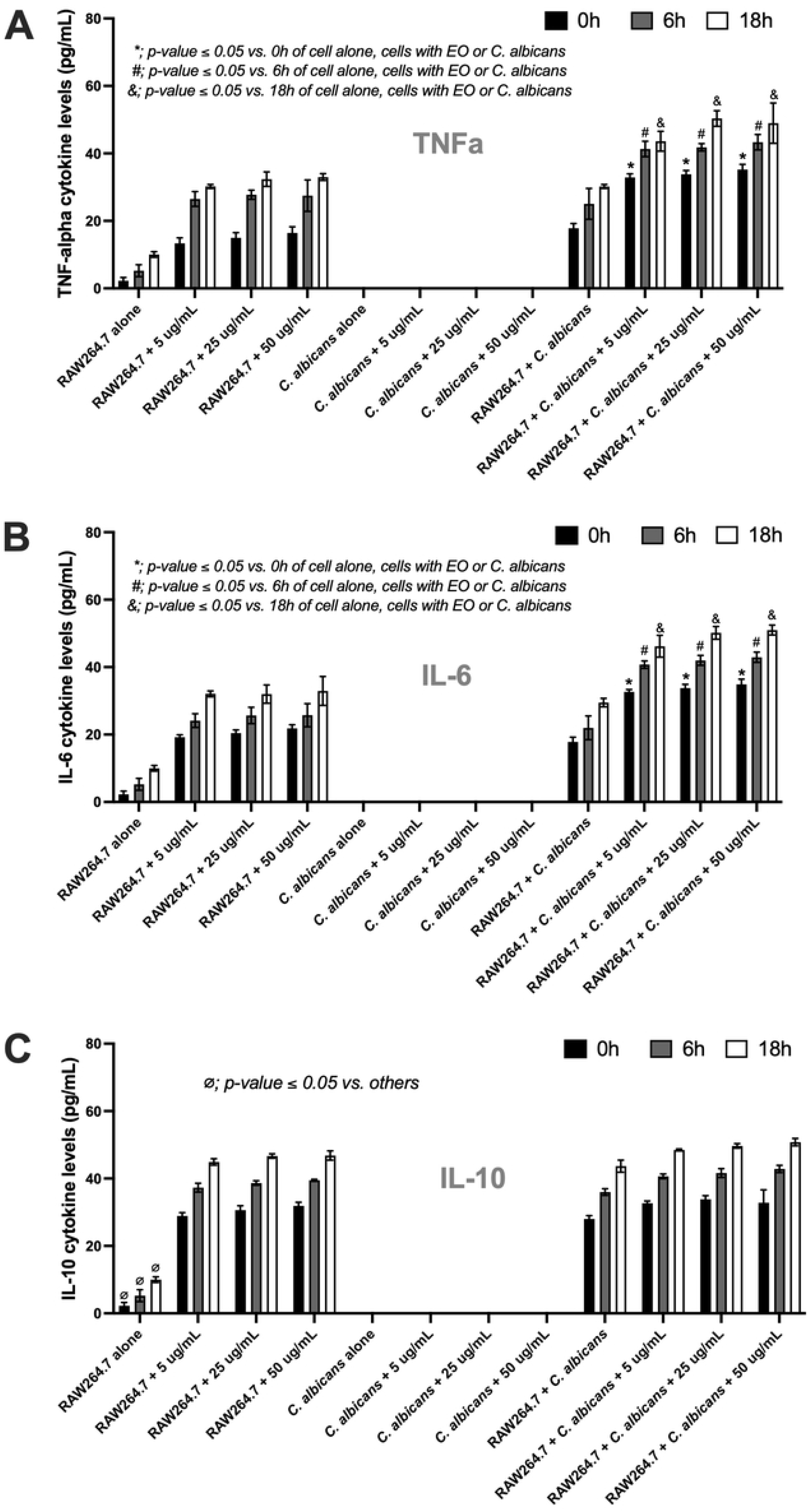
The characteristics of the RAW264.7 macrophage cell line after activation by *C. albicans*, either in the absence or presence of coriander essential oil (0-50 μg/mL), are demonstrated based on cytokine levels of TNF-α (A), IL-6 (B), and IL-10 (C) in the supernatant. Data were collected in triplicate. The data are shown as the mean ± SE, *; *p-value* ≤ *0*.*05* vs. 0 h of other conditions, #; *p-value* ≤ *0*.*05* vs. 6 h of other conditions, &; *p-value* ≤ *0*.*05* vs. 18 h of other conditions, and ∅; *p-value* ≤ *0*.*05* vs. others, between the indicated groups using ANOVA with Tukey’s analysis.

Because immune cell activity is regulated through gene expression, the expression of the *NF-κB* gene was analyzed. The results demonstrated the additive effect of coriander leaf essential oil, especially at 18 h after treatment, the 25 μg/mL and 50 μg/mL of coriander leaf essential oil significantly enhanced the function of macrophage cells by activating the *NF-κB* gene expression in comparison with the untreated *C. albicans* control group (Figure 4A). However, no significant differences were observed in gene expression across varying concentrations of coriander leaf essential oil (5–50 μg/mL) or between different exposure times.

**Figure 4.**
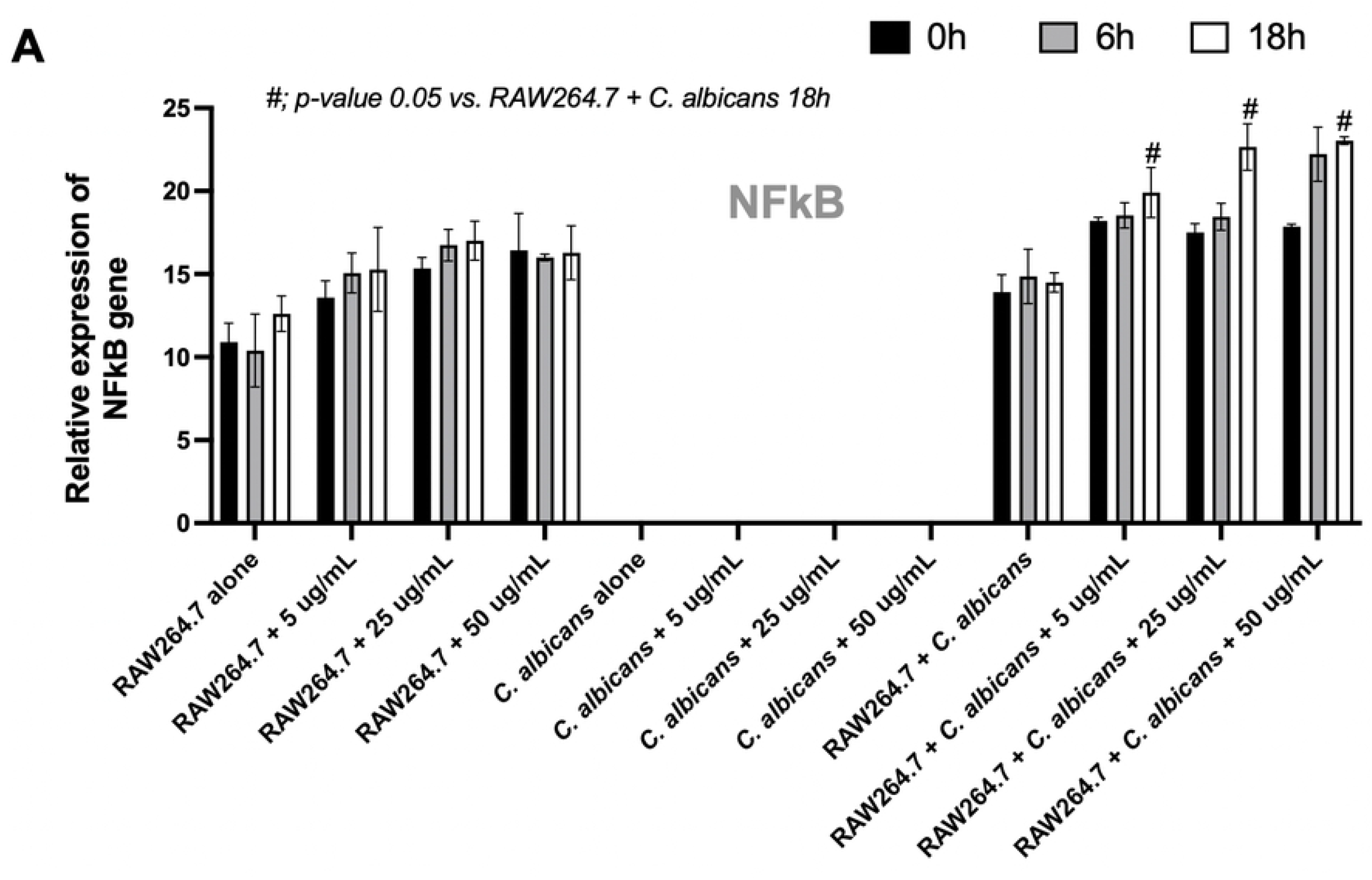
The characteristics of the RAW264.7 macrophage cell line after activation by *C. albicans*, either in the absence or presence of coriander essential oil (0-50 μg/mL), demonstrate the role of inflammatory responses based on *NF-κB* gene expression levels. The experiment was conducted in triplicate. The data are shown as the mean ± SE, #; *p-value* 0.05 vs. RAW264.7 + *C. albicans* 18 h.

## Discussion

In the past decade, essential oils have attracted considerable interest as a rich source of bioactive compounds, with numerous potential health benefits. Essential leaf oil extracted from coriander has been found to contain diverse bioactive components. The major monoterpenoids identified include thymol, carveol, and chrysanthemic acid, all of which have demonstrated antimicrobial and antioxidant activities (11, 12). Additionally, coriander leaf oil also contains several diterpenoid and triterpenoid compounds, including darutigenol, lagochilin, amyrin, and abietic acid, classified within the labdane and abietane structural groups, which are commonly found in plants (13). While their molecular structures have been characterized, common names and detailed bioactivities for many of these compounds are still lacking in current databases; more studies should be conducted. In terms of lipid composition, the coriander leaf essential oil contains several saturated fatty acids, including lauric acid, capric acid, undecanoic acid, dodecanoic acid, and tridecanoic acid. In addition, hydroxydodecanoic acid, a hydroxylated saturated fatty acid, is also present. Notably, previous studies have reported that saturated fatty acids can modulate dendritic cells through Toll-like receptor 4 (TLR4) (14, 15). Furthermore, oxygenated derivatives of long-chain polyunsaturated fatty acids, such as 11-HETE, 9-HEPE, and 12-OxoETE, are associated with modulating inflammation and host immune response (16, 17). This study demonstrates that coriander leaf essential oil supports macrophages in reducing *C. albicans* colonies, which is associated with previous studies (8, 18). The coriander oil components might bind to ergosterol in the *C. albicans* cell membrane, increasing membrane permeability and causing membrane damage and cell death. Future studies should be conducted. Moreover, a previous report shows that coriander leaf essential oil disrupts *Candida* biofilm and adherence (8). Hence, these findings highlight the direct antifungal potential of coriander leaf essential oil.

Additionally, the coriander oil can modulate host immunity. For example, treatment of murine RAW264.7 macrophages with coriander seed oil significantly modulates immune responses, increases phagocytic and killing activity (19), suggesting an enhanced innate immune response. As same as in this study, coriander leaf essential oil raises macrophage immune responses, as evidenced by the enhanced production of pro-inflammatory cytokines (TNF-α and IL-6). These cytokines play essential roles in promoting immune responses against fungal pathogens, especially *C. albicans* (20, 21). Our findings suggest that an immune upregulation is associated with the activation of the *NF-κB* signaling pathway, a key regulator of inflammatory gene expression (22, 23). Mechanistically, the interaction between pathogen-associated molecular patterns (PAMPs), such as β-glucans from *Candida* cell walls, and pattern recognition receptors (PRRs) on macrophages initiates intracellular signaling cascades (24, 25). Among these receptors, Dectin-1 is a well-characterized C-type lectin receptor that specifically binds to β-1,3-glucans (26). Upon ligand engagement, Dectin-1 initiates downstream signaling through spleen tyrosine kinase (Syk), leading to the activation of the CARD9-BCL10-MALT1 complex (27). Leading to the phosphorylation and nuclear translocation of *NF-κB* subunits, facilitating the transcription of pro-inflammatory cytokine genes, including TNF-α and IL-6 (28, 29). Our data suggest that coriander leaf essential oil enhances this process, either by increasing the sensitivity of macrophages to β-glucan stimulation or by directly modulating intracellular signaling pathways. Previously, bioactive components of coriander leaf essential oil, such as thymol and carveol, have been reported to exert immunomodulatory effects, potentially through transcription factors, *NF-κB* (30). It is reliable that these active compounds act synergistically to promote macrophage activation during fungal encounters. Future studies should investigate the functions of those specific active compounds in modulating immune cell responses during *C. albicans* infection.

Interestingly, despite the enhanced pro-/ anti-inflammatory cytokine productions and gene expression, no significant differences were observed across varying concentrations of coriander leaf essential oil (5–50 μg/mL), nor between different exposure times. These results may indicate a saturation threshold in Dectin-1/*NF-κB* axis activation (29), suggesting that even low doses of coriander essential oil are sufficient to trigger maximal response. Furthermore, the observed immune stimulation might have additive effects when combined with fungal β-glucans (31, 32). Coriander leaf essential oil could function as a priming agent, sensitizing macrophages to fungal PAMPs and thereby amplifying Dectin-1-mediated *NF-κB* activation. Collectively, these findings support the potential of coriander leaf essential oil as a natural immunomodulator that enhances macrophage responses via the *NF-κB* pathway. This mechanism may be relevant not only for fungal clearance but also for broader applications in immunotherapy and inflammation regulation, warranting further *in vivo* investigation to clarify its mechanisms and therapeutic potential.

## Conclusion

This study highlights the potential of coriander leaf essential oil as a natural immunomodulatory agent with antifungal activity against *C. albicans*. The findings demonstrate that coriander leaf oil had an additive effect, which can increase macrophage responses, particularly through modulation of killing activity, cytokine production, and the *NF-κB* gene. These results suggest that coriander essential oil may serve as a promising adjuvant therapy, supporting host immune defense while exerting direct antifungal effects.

